# Autophagy Increases Occludin Levels to Enhance Intestinal Paracellular Tight Junction Barrier

**DOI:** 10.1101/2022.04.11.487876

**Authors:** Kushal Saha, Ashwinkumar Subramenium Ganapathy, Alexandria Wang, Nathan Michael Morris, Eric Suchanec, Gregory Yochum, Walter Koltun, Wei Ding, Meghali Nighot, Thomas Ma, Prashant Nighot

## Abstract

**Background and Aim:** Functional loss of paracellular tight junction (TJ) barrier of the gut epithelium and mutations in autophagy genes are factors potentiating inflammatory bowel disease (**IBD**). Previously we showed the role of autophagy in enhancing the TJ barrier via claudin-2 degradation, however, its role in the regulation of the barrier-forming protein occludin remains unknown. Here, we investigate the role of autophagy in the regulation of occludin and its role in inflammation-mediated TJ barrier loss.

**Methods:** Pharmacological and genetic tools were used to study the effect of autophagy on occludin levels and localization, and the role of the MAPK pathway.

**Results:** Autophagy induction using pharmacological activators and nutrient starvation increased total occludin levels in different epithelial cells. Starvation enriched membrane occludin levels and reduced paracellular inulin flux in Caco-2 cells. Starvation-induced TJ barrier enhancement was contingent on the presence of occludin as *OCLN*^-/-^ nullified its TJ barrier enhancing effect. Autophagy inhibited the constitutive degradation of occludin and protected against inflammation-induced TJ barrier loss. Starvation-induced TJ barrier enhancement was prevented by inhibition of autophagy. Autophagy enhanced the phosphorylation of ERK-1/2. Inhibition of these kinases in Caco-2 cells and human intestinal mucosa inhibited the protective effects of autophagy. *In-vivo*, autophagy induction by rapamycin increased occludin levels in mouse intestines and protected against LPS and TNF-α-induced TJ barrier loss. Additionally, acute *Atg7* knockout in adult mice decreased intestinal occludin levels, increasing baseline colonic TJ-permeability and exacerbating the effect of DSS-induced colitis.

**Conclusion:** Our data suggest a novel role of autophagy in promoting the intestinal TJ barrier by increasing occludin levels in an ERK1/2 MAPK-dependent mechanism.

**Graphical abstract:** 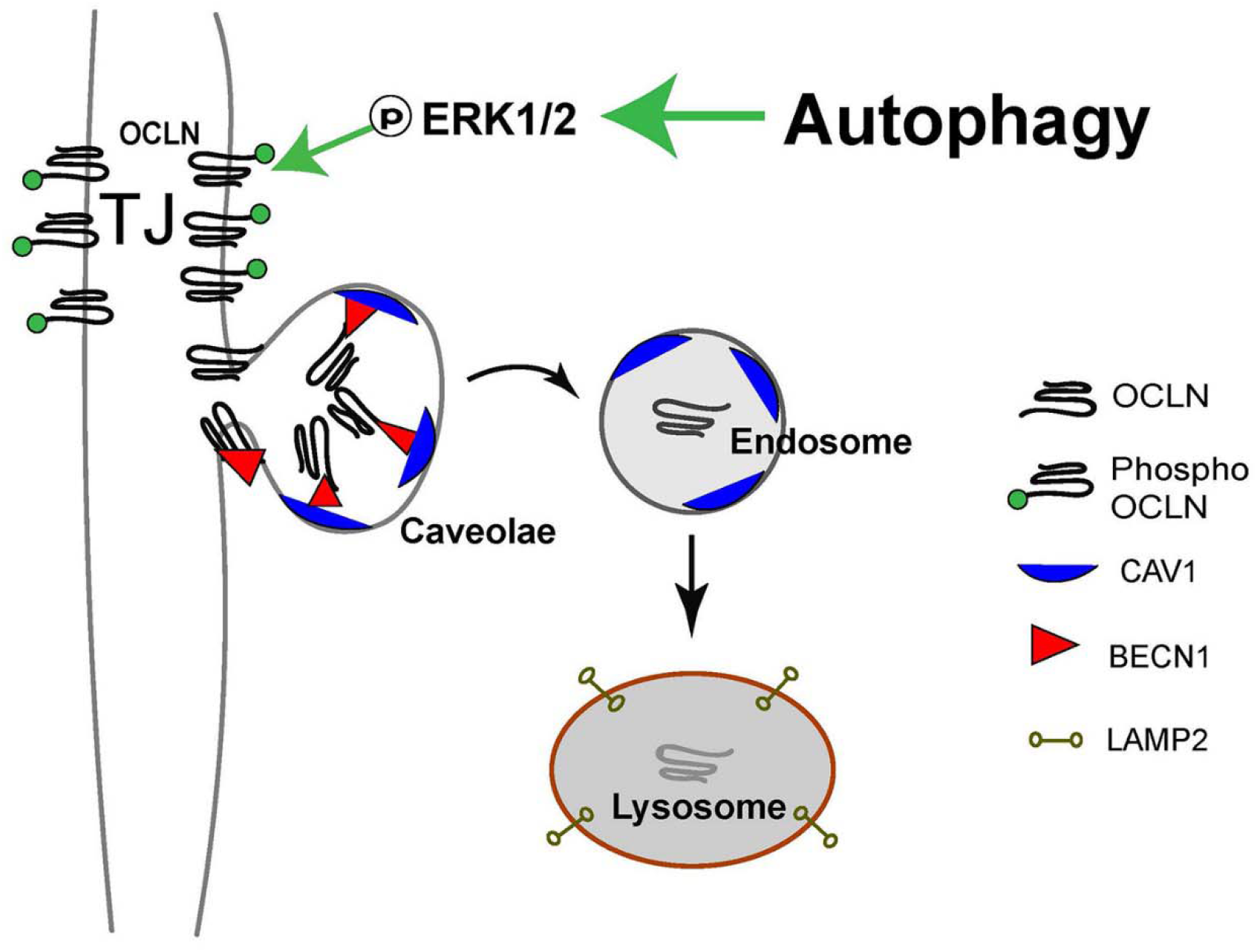

## Introduction

Inflammatory bowel disease (**IBD**), including ulcerative colitis (**UC**) and Crohn’s disease (**CD**), is a chronic inflammatory disorder of the gastrointestinal (**GI**) tract and affects over 1.3% of the adult US (United States) population.^1^ The factors leading to the onset and progression of the disease are multifactorial, however, a key factor attributed to the development, pathogenicity, and severity of this disorder is the increased permeability of the gut epithelium;^2, 3^ compromising the segregation of the luminal contents from the basal vasculature. Paracellular transport is a major route of transport for solutes across the gut epithelium and is regulated through cell-to-cell junctions formed by protein aggregates which seal the intermediate space between adjacent epithelial cells. TJs (Tight Junctions) occupy the apical surface of the lateral plasma membrane and associate with similar TJ plaques on adjacent cells to form a barrier against the free flow of solutes through paracellular space.^4^

Macroautophagy (autophagy) is an evolutionarily conserved mechanism of degradation and recycling of cytosolic contents, serving as a mode of adaptation and protection against cellular stress.^5^ The major steps in autophagy are (i) initiation or formation of the phagophore mediated by two key protein kinase complexes, ULK1/2-ATG13-RB1CC1/FIP200 complex and the BECN1/beclin1-PIK3C3/Vps34-PIK3R4/Vps15-ATG14 complex; (ii) expansion of the phagophore by ubiquitin-like proteins ATG12 and GABARAP/LC3; (iii) autophagosome formation with LC3 modification (lipidation of cytosolic LC3-I to LC3-II) and binding to autophagosome membrane; and (iv) the fusion of the autophagosome with the lysosome to form an autolysosome which degrades the cargo proteins.^6^ Studies have linked defects in autophagy to IBD development, however, a mechanistic understanding of the role of autophagy in the regulation of the gut epithelial barrier and the basis of defective autophagy in IBD development remains poorly understood. Being an intracellular degradation pathway, autophagy intersects with endocytic processes in a tightly controlled manner.^7–9^ In our previous work, we demonstrated the autophagy pathway’s TJ barrier protective role through the clathrin-mediated degradation of the pore-forming protein claudin-2.^10, 11^

Occludin is a MARVEL (MAL-related proteins for vesicle trafficking and membrane link) domain-containing protein family member. It is a tetra spanning protein with two extracellular loops, one intracellular loop, a large cytoplasmic C-terminal domain, and a short N-terminal domain. Occludin interacts with the guanylate kinase domain of ZO-1 via the coiled-coil region of the C-terminal domain.^12^ While several studies have highlighted the TJ barrier protective role of this protein against the flow of macromolecules, the precise mechanism of how occludin participates in this process remains unknown. In addition to the barrier-enhancing role of occludin, loss of this protein and depleted paracellular barrier are hallmarks of IBD. In this study, we investigated the effect of autophagy on occludin expression and the TJ barrier against macromolecule flux. Here we report a novel role of autophagy in the upregulation of occludin protein levels and enhancement of the TJ barrier against macromolecular flux. We also demonstrate the protective effect of autophagy and occludin up-regulation against inflammatory cytokine-induced loss of the paracellular TJ barrier and highlight the role of the Mitogen-Activated Protein Kinase (MAPK) proteins ERK-1/2 in this process.

## Materials and Methods

### Chemicals and antibodies

Rapamycin was purchased from Life Technologies (PH21235), cycloheximide (C7698), LPS O127:B8 (L3129), SBI-0206965 (SML1540), iodoacetamide (I1149) were obtained from Sigma. Murine TNF-α (50349-MNAE) was purchased from SinoBiologicals (Wayne, PA, USA). Human IFN-γ (300-02) and TNF-α (300-01A) were purchased from PeproTech and IL17A (ILA-H5118) from Acrobiosystems. Bafilomycin A1 was purchased from Santa Cruz Biotechnology, (sc-201550), U0126 from TORCS (1144), and PD98059 from Calbiochem (513000). [^3^H] Inulin (specific activity 100 mCi/ mmol) was purchased from American Radiolabelled Chemicals (ART117). 2-Mercaptoethanesulphonic Acid (MESNA) was purchased from Acros Organics (443150250). The primary antibodies used included anti-occludin, anti-caveolin-1, anti-ATG7, anti-SQSTM1/p62, anti-β-actin (ProteinTech, 27260-1-AP, 66067-1-1g, 26523-1-AP, 27355-1-AP, 10088-2-AP, 19812-1-AP, 18420-1-AP, HRP-60008, respectively), anti-beclin-1(Abcam 207612), anti-Y14-phospho caveolin-1, anti-phospho Threonine, anti-Phospho Tyrosine, anti-ERK-1/2, Phospho-p44/42 MAPK (Erk1/2) (Thr202/Tyr204) and anti-LAMP2 (Cell Signaling Technologies 3251S, 9386S, 9411S, 4695S, 4377S and 4906S respectively). The anti-rabbit HRP, anti-mouse HRP, secondary antibodies, and all other molecular biology grade reagents were purchased from various commercial vendors. TaqMan RT primers against human occludin (Hs05465837_g1) and GAPDH (Hs02786624_g1) were purchased from Invitrogen.

### Cell Culture

Human intestinal epithelial Caco-2 cells and T84 cells (ATCC) and MDCK-II cells (ECACC) were maintained in Dulbecco’s Modified Eagle’s Medium (DMEM) – High Glucose (Gibco, Cat. No. 11965118) supplemented with 10% heat-inactivated fetal bovine serum and antibiotics at 37°C in a 5% CO_2_ incubator. Caco-2 cells were grown on 0.4 μm pore size, 12-mm diameter filter inserts. The transepithelial electrical resistance (TER) of cells was measured by an epithelial voltohmeter (World Precision Instruments, Sarasota, FL). Monolayers with a TER of 450–500 Ω/cm^2^ were used for experiments. Nutrient starvation was induced by replacing DMEM with serum-free Earle’s Balanced Salt Solution (EBSS) (Sigma, E3024).

### Determination of Caco-2 Paracellular Flux

Permeability was determined using the paracellular marker inulin (^3^H, M_r_ = 5000). The apical-to-basal flux of inulin was determined by adding it to the apical solution. Radioactivity was measured in the basal solution at 30 min and 60 min using a scintillation counter, as described previously in.^13^

### Western Blot Analysis

Western blot analysis was performed by electrophoresing protein samples in a 4 %-15% SDS-PAGE gel as discussed by us previously.^9^

### Transmission electron microscopy

Caco-2 cell monolayers grown on a 0.4 μm membrane were subjected to starvation as mentioned above. As previously described, the control and starvation-induced samples were prepared for immune-gold staining and TEM.^11^

### Confocal immunofluorescence

Confocal Immunofluorescence for occludin, ERK1/2, caveolin-1, and beclin-1 was performed by standard methods discussed previously. The slides were examined using a confocal fluorescence microscope Leica SP8. Images were processed with LAS X software (Leica Microsystems). Images are presented as maximum intensity projections rendered from 30 z-stacks of 0.30 μm. Representation of several fields from at least 3 separate samples.

### Co-immunoprecipitation

Using methods previously discussed, the co-immunoprecipitation experiments were performed using protein G dynabeads from Invitrogen (10004D) according to the manufacturer’s protocol.^11^

### Quantitative Real-Time PCR

Cells were then dissolved in TRIZol (Invitrogen, 15596026) followed by RNA isolation using the Direct-zol RNA Miniprep Plus kit (Zymo Research, R2072) by following the manufacturer’s instructions. Transcript levels of occludin and GAPDH were quantified as discussed previously.^14^

### Cell surface biotinylation and endocytosis assay

Surface proteins on Caco-2 cells were biotinylated using Pierce Cell Surface protein biotinylation and Isolation Kit (Thermo Scientific, A44390) following the manufacturer’s instructions. Cells were then incubated in either normal media or EBSS at 37°C for 0, 3, or 6 hours. Isolation and determination of biotinylated protein fractions were performed as previously described.^13^

### CRISPR/Cas9 knockout of occludin, ATG7, ERK-1, and ERK-2

Single guide RNA (sgRNA) targeting the region TGAGCAGCCCCCCAATGTCG of *OCLN* or AAATAATGGCGGCAGCTACG of *ATG7* or CGGGGAGCCCCGTAGAACC region of ERK-1 (*MAPK-3*) or CGCGGGCAGGTGTTCGACGT region of ERK-2 (*MAPK-1*) or scrambled sgRNA for control in pCRISPR-LVSG03 (Genecopoeia) was used to generate *OCLN*-/-, MAPK *dKO*, *ATG7*^-/-^ and Scr Caco-2 cells as discussed previously.^11^ Occludin ORF in pCMV6-AC-GFP (Origene) and corresponding control plasmid were used to transfect Caco-2 cells using Lipofectamine 2000 (Invitrogen, 11668027) as per the manufacturer’s instructions.

### Experimental animals

Experimental methodologies used in the study were approved by the Institutional Animal Care and Use Committee (IACUC) of The Pennsylvania State University College of Medicine. Adult mice were engineered with floxed alleles of *Atg7* and a transgene expressing the TAM-regulated Cre recombinase fusion protein under the control of the ubiquitously expressed ubiquitin C (UBC) promoter.^15^ Atg7 deficiency was created by providing tamoxifen (Sigma Cat No. T5648; 20 mg/ml suspended in 98% sunflower seed oil and 2% ethanol mixture) to 10-week-old mice as described previously.^11^ The wildtype C57BL/6J mice (Stock No. 000664, Jackson Laboratory) were treated with rapamycin (40 mg/kg/day for 3 days, i.p injection). In the dextran sulfate sodium (DSS) model, mice were given 2.5% DSS in drinking water for 7-days, as detailed previously.^14^ LPS and TNF-α (0.2μg/kg each) were dissolved in sterile PBS and injected i.p followed by incubation for 12 and 4 hours respectively.

### Measurement of paracellular permeability of murine and human colonic tissues

The paracellular permeability of the mice and human intestinal tissues was measured using 0.03-cm^2^-aperture Ussing chambers (Physiologic Instruments, CA) and mucosal-to-serosal flux of [^3^H]-inulin, as described by us previously.^16^ Transepithelial electrical resistance (TER, Ω·cm^2^) was calculated from the spontaneous potential difference and short-circuit current.

### Human Tissue samples and treatment

The surgically resected fresh human intestinal tissue samples were obtained from the Department of Surgery, Division of Colon and Rectal Surgery as per the protocols approved by the Institutional Review Board. Tissues were ascribed as normal or diseased by the Department of Pathology and Laboratory medicine, Penn State College of Medicine. Tissues were processed as described previously^11^ and incubated in the presence and absence of 300 nM Rapamycin (Alfa Aesar, J62473) with or without 25 μM U0126 or 20 μM PD98059.

### Statistical analysis

Data are reported as means ± SEM. Data were analyzed by using appropriate statistical tests (Sigma Stat, Systat Software, San Jose, CA).

## Results

### Autophagy increases cellular occludin and its membrane localization causing decreased macromolecular flux across the paracellular space

We have previously shown that autophagy enhances the intestinal TJ barrier by reducing the levels of the pore-forming TJ protein claudin-2 through its interaction with the μ subunit of the clathrin adaptor protein AP2 and autophagy receptor LC3.^10, 11^ Besides claudin-2, the barrier-forming protein occludin is another major component of the TJ barrier that regulates macromolecular flow across the paracellular space. To examine if autophagy affects occludin, we examined occludin levels in Caco-2 cells upon starvation and found a significant increase in occludin protein levels (**Fig. 1A and B**). We further assessed the effect of starvation on other epithelial cell lines like MDCK-II and T84. Like Caco-2 cells, starvation increased occludin levels in both cell lines (**Fig. 1C**), indicating that the starvation-induced increase in occludin protein is not exclusive to the Caco-2 cells. Next, we examined the effect of known chemical inducers of autophagy on cellular occludin levels. Rapamycin^10^, resveratrol^17^, SMER28^18^, and Metformin^19^ treatment induced autophagy as evident by a reduction in autophagy substrate p62/SQSTM1 and significantly increased occludin protein levels in Caco-2 cells (**Fig. 1D and E**). Overall, the data shows that induction of autophagy by different mechanisms increases cellular occludin levels. Next, we investigated the fate of this increased cellular occludin. For this, we performed a fractionation analysis to determine the membrane and cytosolic distribution of occludin in Caco-2 cells upon starvation. We found that starvation enhanced the levels of occludin in the membrane fractions compared to control cells (**Fig. 1F and G**). Furthermore, confocal immunofluorescence imaging (**Fig. 1H, I**) and transmission electron microscopy (TEM) (**Fig. S1A**) also showed that starvation enhanced the localization of occludin to the TJs in Caco-2 cells and mouse colon-derived organoids (**Fig. S1B**). In addition, we found occludin immunoprecipitates from Caco-2 cells to have significantly increased Thr phosphorylation and a corresponding reduction of Tyr phosphorylation upon starvation (**Fig. S1C**).^20^

**Figure 1.**
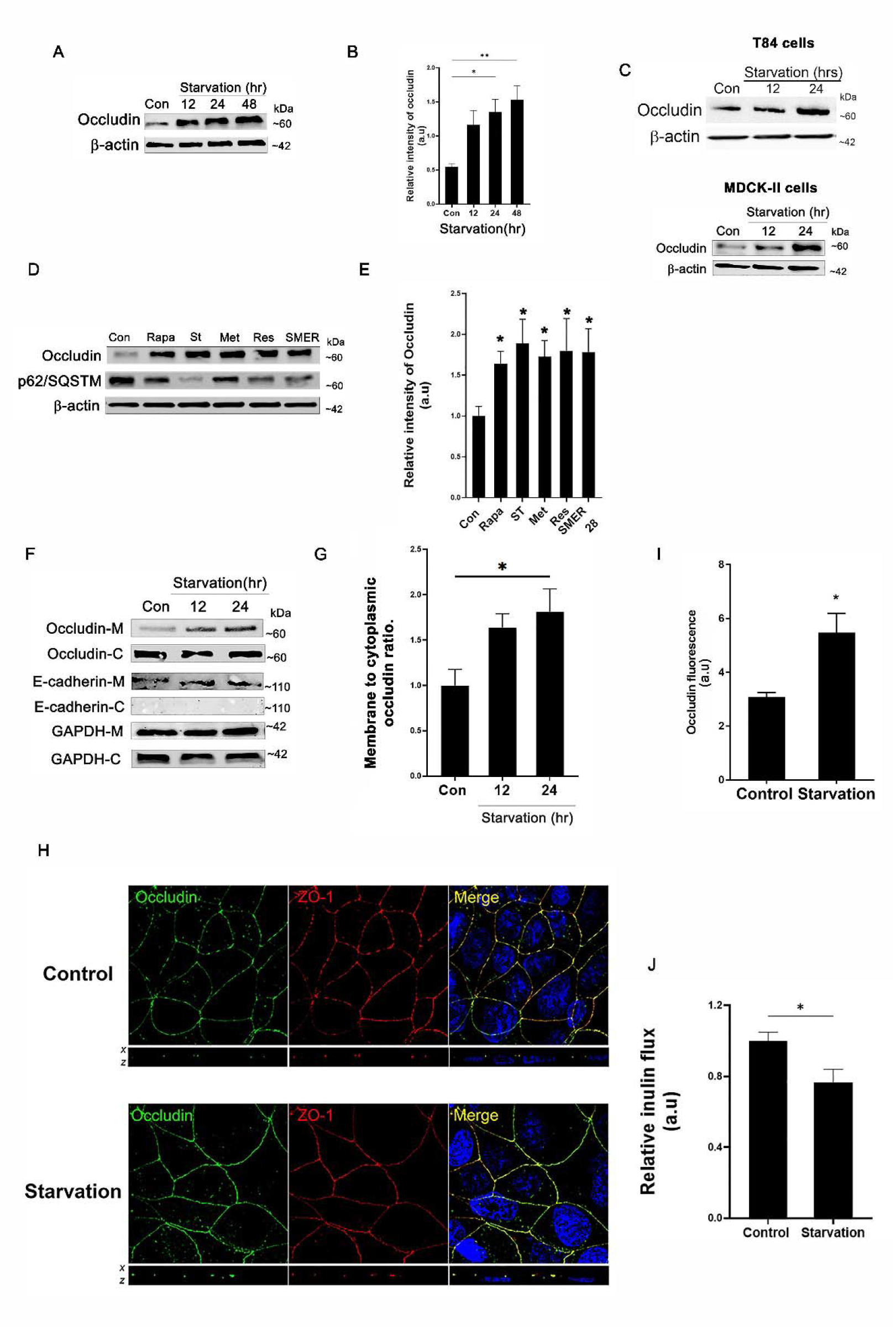
Autophagy enhances cellular occludin levels and enhances the TJ barrier against macromolecule flux. **(A)** Western blot for occludin from control and 12 and 24 hr starvation-induced Caco-2 cells. β-actin is shown as a loading control. **(B)** Quantification of occludin in panel A using ImageJ software to indicate the relative levels of occludin after starvation. n > 3 independent experiments. **(C)** Western blot for occludin from control and starvation-induced MDCK-II and T84 cells. β-actin is shown as a loading control. Like Caco-2 cells, starvation increased occludin levels in both cell lines. Blot represents > 3 independent experiments. **(D)** Western blot for occludin from Caco-2 cells treated with rapamycin (Rapa, 500nM), small-molecule enhancer 28 (SMER, 50μM), resveratrol (RES, 100μM), metformin (MET, 100μM) and EBSS. SQSTM/p62 protein degradation is shown as evidence for autophagy induction in the different treatments. β-actin is shown as a loading control. The protein levels of occludin increased by all the autophagy inducers. **(E)** Quantification of occludin levels in panel D. n ≥ 3 independent experiments. **(F)** Starvation increased the relative presence of occludin in the membrane fraction. M: membrane; C: cytoplasm. E-cadherin is shown as a marker for cell membrane and cytoplasmic fractions. GAPDH is shown as a loading control for membrane fractions. **(G)** quantification of the membrane to cytoplasmic occludin ratio. The blots are representative of ≥ 3 independent experiments. **(H)** Starvation increased the presence of occludin (green) on the cell membrane and increased co-localization with the TJ protein ZO-1 (red). nuclei (blue). White bar=5 μm. **(I)** Quantification of occludin fluorescence intensity from panel H. **(J)** Starvation significantly reduced the flux of inulin across confluent Caco-2 cell monolayers. n > 5 independent experiments. Statistical analysis by Student’s *t-test* or one-way ANOVA with Tukey’s post-test. *, *p* < 0.05; **, *p* < 0.01.

We have previously shown that starvation increases transepithelial resistance (TER) and decreases the paracellular flux of the small solute urea, which is dependent on claudin-2 degradation.^10^ Occludin is, however, known to regulate the transport of larger, uncharged solutes across the paracellular space otherwise referred to as the leak pathway.^21^ To test whether starvation has a barrier-enhancing effect against the leak pathway, we assessed the paracellular flux of inulin across Caco-2 cell monolayers. We observed that starvation significantly decreased the paracellular flux of inulin (**Fig. 1J**). To assess if this starvation-induced reduction in inulin flux is caused by the autophagy-mediated degradation of claudin-2, we quantified the inulin flux across control and starved *CLDN2*^-/-^ Caco-2 cell monolayers and observed no significant difference (**Fig. S1D).** Taken together, these studies show that starvation increases the levels of TJ localization of the occludin protein and enhances the TJ barrier against the flow of the large solute inulin.

### Autophagy reduces occludin endocytosis and enhances the protein half-life

We next examined the possible mechanism underlying the starvation-mediated upregulation of occludin. We observed no significant differences in occludin transcript levels in Caco-2 cells upon starvation (**Fig. S2A)**. Thus, we investigated the possibility of reduced occludin protein degradation. For this, control and pre-starved Caco-2 cells were treated with the protein synthesis inhibitor cycloheximide for increasing time points. Western blot analysis showed a temporal reduction of the occludin protein in the control samples. In contrast, occludin protein levels in the pre-starved Caco-2 cell samples showed no significant change, suggesting that starvation protects occludin from degradation (**Fig. 2A**). To further investigate the starvation-mediated enhancement of membrane occludin, we assessed the effect of starvation on occludin endocytosis. Previous studies have shown that occludin undergoes caveolar endocytosis by interacting with caveolin-1, the primary protein component of caveolar structures. ^13, 22^. Additionally, we have previously reported the autophagy-independent role of the Atg6/Beclin-1 protein in the constitutive caveolar endocytosis and lysosomal degradation of occludin^23^. Therefore, we probed occludin immunoprecipitates from control and starved Caco-2 cells for caveolin-1, beclin-1, and the lysosomal marker LAMP-2. The Co-IP showed a temporal reduction in the interaction of occludin with all three proteins upon starvation (**Fig. 2B and C**). We also assessed if starvation affects the global caveolae-mediated endocytosis machinery and determined the levels of pY-14-Cav-1 at different time points of starvation. Our data shows that with increasing time points of starvation the levels of pY-14-Cav-1 increased, indicating that the caveolae-mediated endocytosis process remains unaffected during autophagy (**Fig. S2B**) ^24, 25^. We also assessed the endocytosis of cholera toxin-β-subunit in control and starved Caco-2 cells as a known marker for caveolar endocytosis.^26^ Confocal microscopy showed no significant difference in the ability of Caco-2 cells to endocytose the toxin upon starvation further indicating that starvation does not inhibit the overall caveolar endocytosis pathway (**Fig. S2C**).

**Figure 2.**
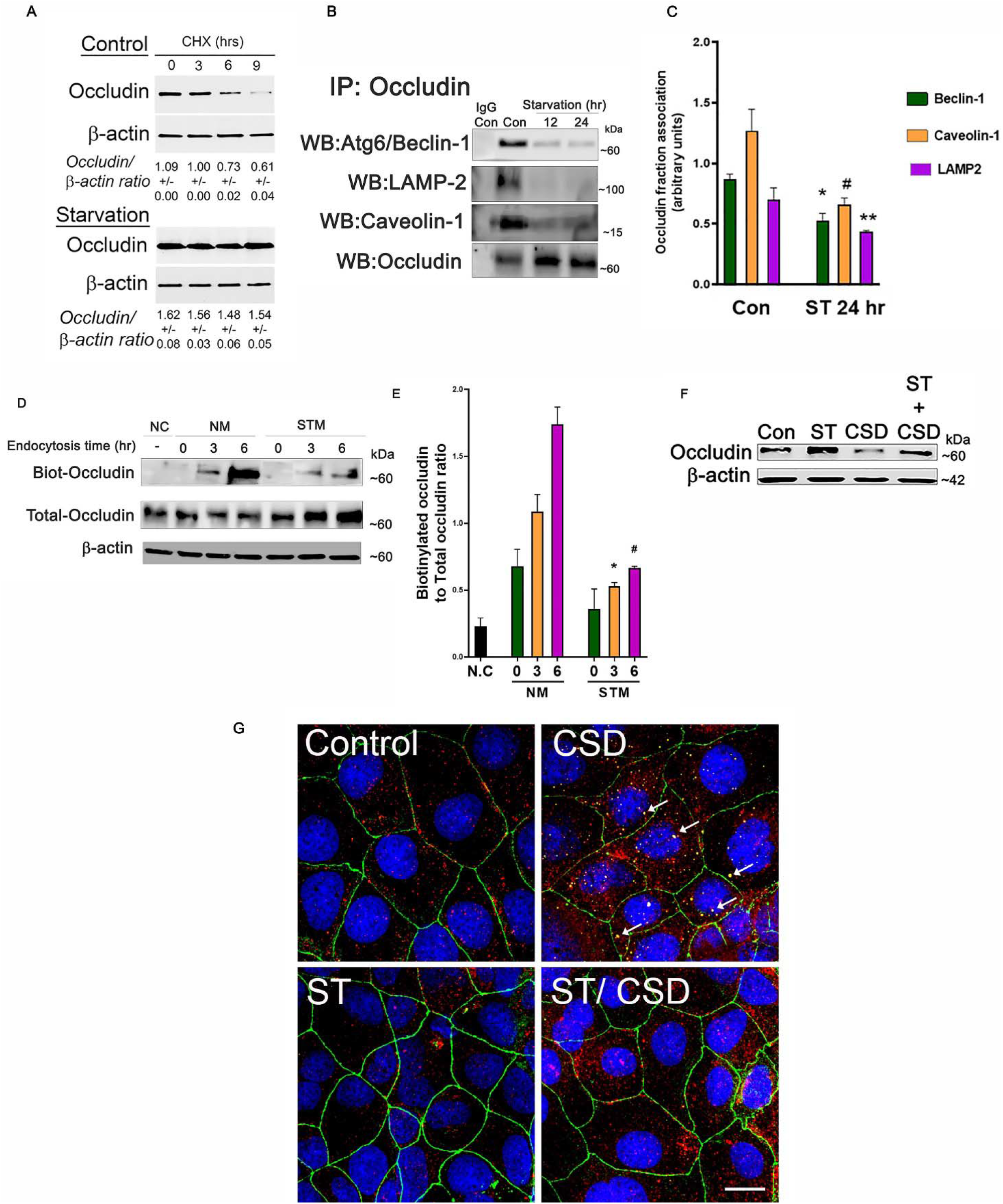
Autophagy enhances the half-life of occludin protein and protects it from constitutive degradation. **(A)** Cycloheximide treatment did not affect occludin levels upon starvation. β-actin is shown as a loading control. Quantification of relative occludin levels is denoted below the lanes as mean ± SEM. n > 3 independent experiments. **(B)** Co-immunoprecipitation studies showed decreased association of occludin with beclin-1, caveolin-1, and lysosomal marker LAMP-2 from 12 hours post starvation. The negative control includes immunoprecipitation with control IgG. **(C)** Quantification of beclin-1, caveolin-1, and LAMP2 observed in occludin immunoprecipitations from panel B. n = 3. **(D**) Starvation reduces the endocytosis of membrane-bound biotinylated occludin. **(E)** Quantification biotin-labeled to total occludin ratio from panel D. n = 3. **(F)** Starvation protects occludin from caveolin-1 scaffolding domain (CSD) peptide (5μM)-induced occludin degradation. n ≥ 3 independent experiments. **(G)** Starvation reduced the CSD-induced co-localization between occludin (green) and caveolin-1 (red). White arrows denote caveolin-1 and occludin co-localization (yellow color). Data representative of > 3 independent experiments. White bar=5 μm. Statistical analysis by Student’s *t-test* or one-way ANOVA with Tukey’s post-test. *, # and **, *p* < 0.01 *vs.* each other.

We next directly assessed the effect of starvation on occludin endocytosis using biotin labeling.^13, 27^ For this, following biotinylation of membrane proteins, the Caco-2 cells were subjected to starvation for different periods followed by removal of all surface biotin and isolation of endocytosed biotinylated proteins. We observed increased levels of biotinylated occludin in control cell lysates compared to starved cells, indicating reduced endocytosis of occludin after starvation (**Fig. 2D and E**). Additionally, we tested the effect of Caveolin-1 scaffold domain peptide (CSD) treatment on starvation-induced Caco-2 cells. We have previously shown that treating Caco-2 cells with this peptide induces occludin degradation by enhancing its endocytosis in caveolar pits.^23^ Assessment of occludin levels in control and pre-starved Caco-2 cells showed a very significant reduction in occludin levels upon CSD treatment alone, however, starvation prevented CSD-mediated reduction in occludin levels (**Fig. 2F).** Furthermore, confocal imaging of these samples showed increased co-localization between occludin and caveolin-1 upon CSD treatment which was absent in the cells subjected to pre-starvation (**Fig. 2G**). Taken together our data shows that starvation protects occludin from endocytosis and degradation by disrupting its association with caveolin-1 without compromising the overall caveolae-mediated endocytosis machinery.

### Starvation-induced enhancement of the TJ barrier is autophagy and occludin dependent

We next assessed whether autophagy is essential for the starvation-induced upregulation in cellular occludin levels and the TJ barrier enhancement. For this, Caco-2 cells were subjected to starvation in the presence and absence of the autophagy inhibitors bafilomycin A1 or SBI-0206965.^11, 28^ We found that both autophagy inhibitors prevented the starvation-induced increase in the occludin levels (**Fig. 3A and B**). Additionally, these inhibitors nullified the effect of starvation on the TJ barrier by preventing the starvation-induced reduction in the paracellular inulin flux (**Figure 3 C**). To further verify the role of autophagy in TJ barrier enhancement and occludin upregulation, and to negate the non-specific, off-target effects of these inhibitors, we used CRISPR/Cas9 to generate *ATG7* knockout (*ATG7*^-/-^) Caco-2 cells.^29^ We have previously successfully demonstrated that deletion of the *ATG7* gene in Caco-2 cells disrupts the autophagy pathway.^11^ To this end, we observed no changes in occludin levels upon starvation in *ATG7*^-/-^ cells compared to Caco-2 and non-target guide RNA transfected Scr cells (**Fig. 3D and E**). Additionally, unlike Scr cells, starvation in *ATG7*^-/-^ cells did not reduce the inulin flux (**Fig. 3F**) and the interaction between occludin and caveolin-1 upon CSD treatment (**Fig. 3G; S3A and S3B).** Taken together these data demonstrate the importance of the autophagy pathway for the upregulation of occludin protein levels and enhancement of the paracellular barrier against the flow of macromolecules.

**Figure 3.**
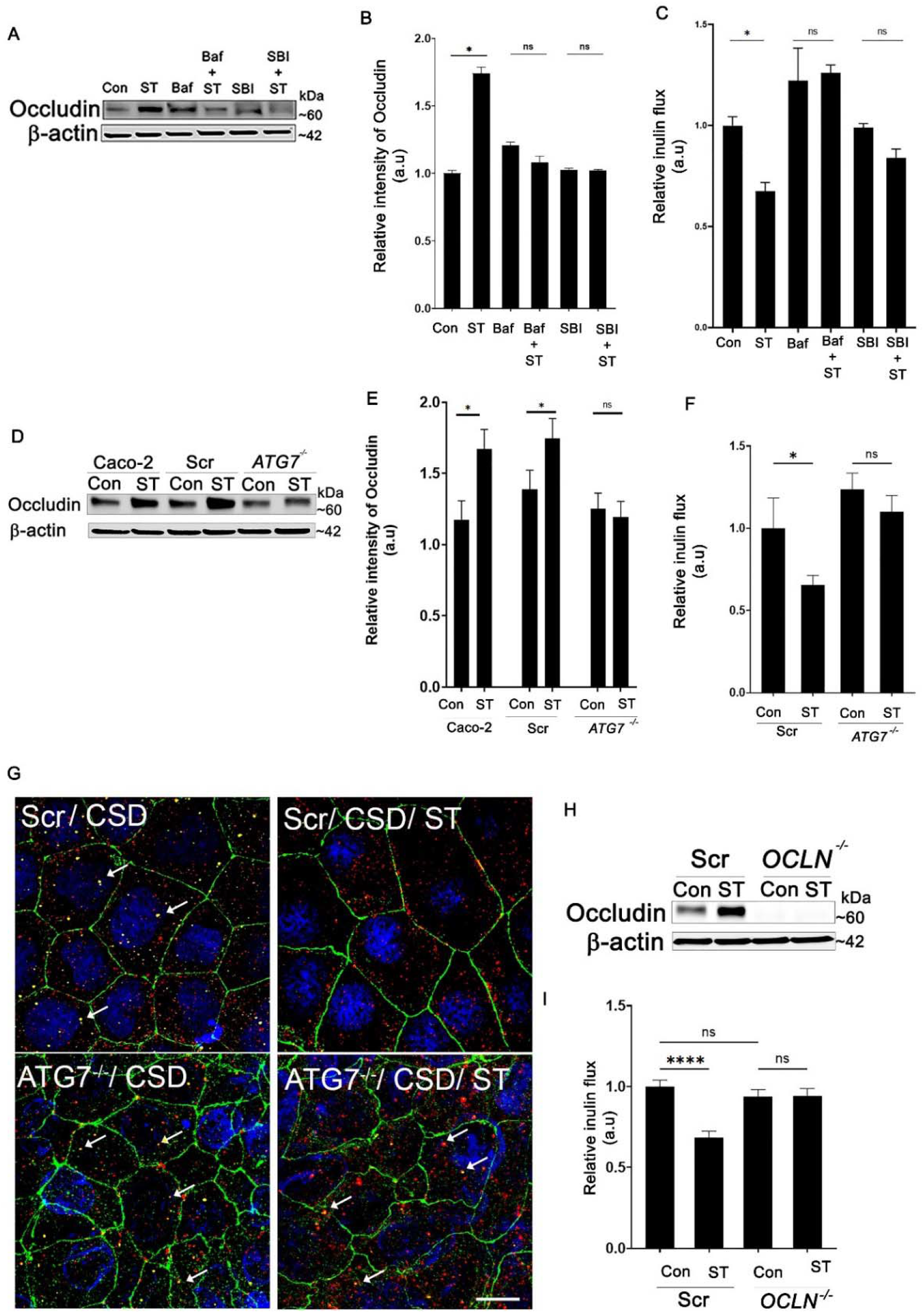
Autophagy and occludin are essential for starvation-induced enhancement of the TJ barrier. **(A)** Autophagy inhibitors bafilomycin A (*BAF*, 20 nM) or SBI-0206965 (SBI, 30 μM) prevented the starvation-induced increase in occludin levels in Caco-2 cell monolayers. **(B)** Quantification of occludin protein levels in panel A. **(C)** BAF and SBI also prevented the starvation-induced decrease in the flux of inulin **(D)** The autophagy-deficient ATG7^-/-^ failed to upregulate levels of cellular occludin upon starvation, unlike Caco-2 and non-target sgRNA transfected Caco-2 (Scr) cells. **(E)** Quantification of occludin protein panel D. **(F)** The ATG7^-/-^ cells failed to reduce the paracellular flux of inulin upon starvation. **(H)** Starvation did not prevent the co-localization between occludin (green) and caveolin-1 (red) upon CSD treatment in the ATG7^-/-^ cells. White bar = 5 μm. White arrows denote occludin-caveolin-1 co-localization. **(H)** Western blot showing the efficiency of CRISPR/Cas9 mediated knockout of occludin in Caco-2 cells. **(I)** Inulin flux remained unchanged upon starvation in the *OCLN*^-/-^ cells. n > 6 independent experiments. Statistical analysis by Student’s *t-test* or one-way ANOVA with Tukey’s post-test. *, *p* < 0.05; ****, *p* < 0.0001; ns – non-significant.

We next tested whether occludin is needed for the starvation-induced TJ barrier enhancement against macromolecular flux. For this, we utilized CRISPR/Cas9 to generate occludin knockout (*OCLN*^-/-^) Caco-2 cells. Transfection of Caco-2 cells with guide RNA targeting occludin resulted in the complete knockout of occludin (**Fig. 3H**). We observed no significant difference between the baseline inulin flux between the Scr and *OCLN* cells, however, while the inulin flux was significantly reduced in the Scr cells upon starvation, there was no significant change in inulin flux in the *OCLN*^-/-^ cells (**Fig. 3I**). Additionally, no significant difference in the baseline TER between the Scr and *OCLN*^-/-^ cells were observed and starvation significantly increased the TER of both cell types (**Fig. S3C)**. Taken together these data demonstrates the need for occludin in the autophagy-induced reduction of paracellular macromolecule permeability.

### Starvation-induced enhancement of cellular occludin levels and TJ barrier is mediated by ERK-1/2

Previous studies have shown the critical role of the Mitogen-Activated Protein Kinase (MAPK) pathway in both autophagy^30^ and membrane localization of TJ proteins.^31^ Adding to these reports, our analysis of the Caco-2 proteome indicates association between the ERK1/2 kinases, occludin, autophagy, and associated endocytic components (**Fig. S4A)**. Therefore, we tested the role of these kinases in the starvation-induced enhancement of the TJ barrier. We observed that starvation activated the MAPK proteins ERK-1 and 2 as evidenced by the increased phosphorylation of these kinases (**Fig. 4 A and B**). Pharmacological inhibition of the kinases with either U0126 or PD98059 prevented the starvation-induced reduction of inulin flux (**Fig. 4C**) and the starvation-induced increase in occludin localization to the membrane (**Fig. S4B, S4C**). In contrast, p38 or JNK MAPK inhibition did not impact the autophagy-induced reduction in macromolecular flux (data not shown). We next sought to assess the effect of genetic deletion of ERK1/2 on starvation-induced phenotypes. For this, we employed CRISPR/Cas9 to knock out the ERK-1 and 2 genes in Caco-2 cells (*dKO*) (**Fig. 4D**). We observed no significant changes in baseline inulin flux between Scr and dKO cells (data not shown). Additionally, like the non-target Scr cells, the TER significantly increased in the *dKO* cells upon starvation (**Fig. S4D**), however, starvation failed to reduce the flux of inulin in the *dKO* cells (**Fig. 4E).** Since TER is a metric used to determine ionic and small solute paracellular permeability across the pore pathway, this observation supports the presence of distinct mechanisms of regulation for the solute transport across paracellular space through the pore and leak pathways. In addition to the effect of ERK1/2 deletion on the TJ barrier, we assessed the effect of the absence of the ERK1/2 kinases on occludin endocytosis. Probing the occludin immunoprecipitates of *dKO* cells showed that starvation failed to reduce the association of occludin with caveolin-1, beclin-1, and LAMP-2 (**Fig. 4F and G**). Furthermore, in the *dKO* cells, starvation had no effect in preventing the association between occludin and caveolin-1 upon CSD treatment (**Fig 4H; S4E and S4F)**. Taken together these studies highlight the essential role of the ERK-1/2 kinases in protecting occludin from degradation and in regulating preventing macromolecular flux across the leak pathway upon autophagy induction.

**Figure 4.**
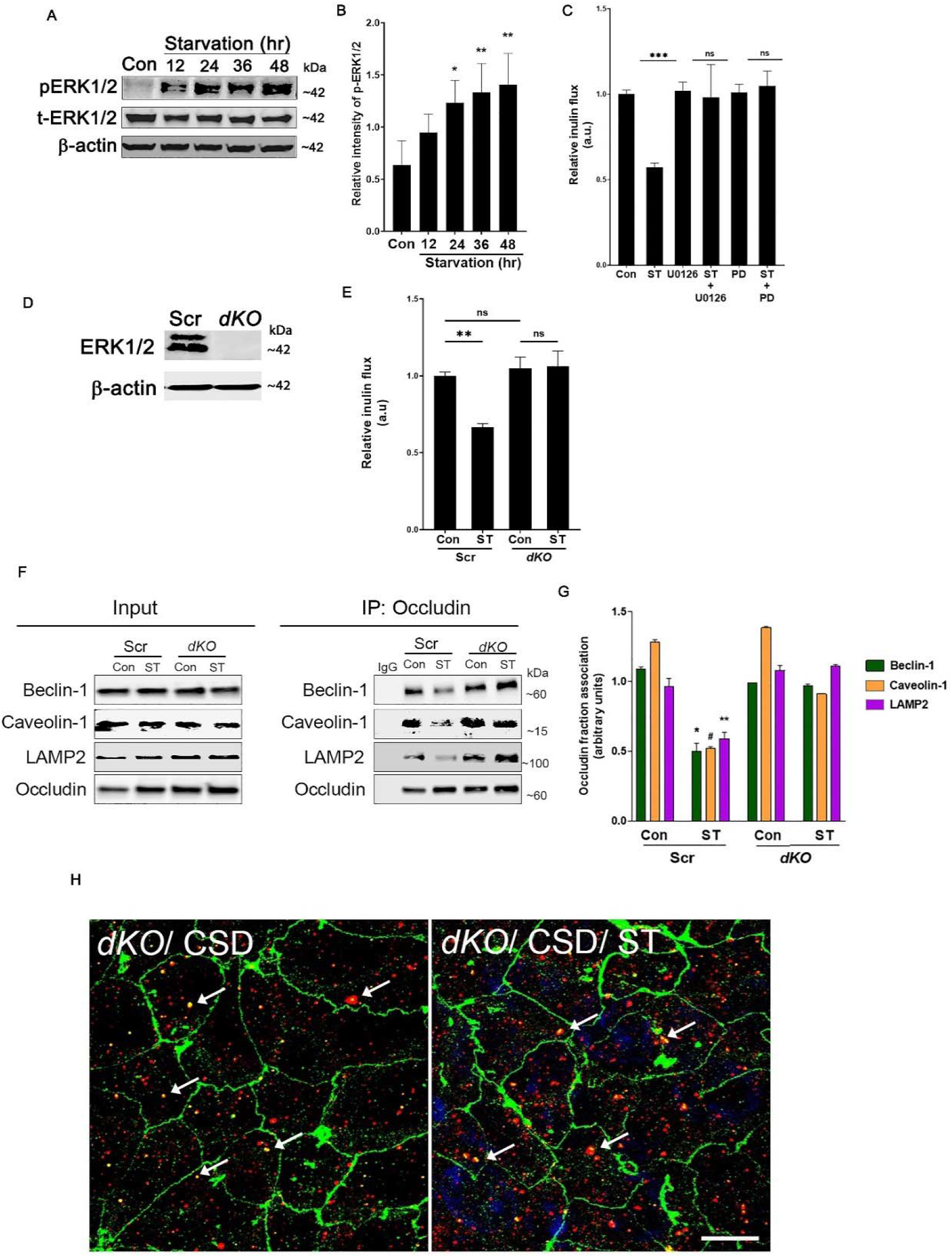
The MAPK proteins ERK-1 and ERK-2 are essential for the autophagy-induced upregulation of occludin. (A) ERK1/2 phosphorylation increased after starvation. β-actin is shown as a loading control. The blot is representative of 3 independent experiments. **(B)** Quantification of phospho-ERK1/2 levels in panel A. **(C)** Inhibition of ERK1/2 activation with U0126 (25 μM) or PD98059 (20 μM) treatment prevented the starvation-induced reduction in inulin flux. **(D)** Western blot showing the efficacy of CRISPR/Cas9 mediated knockout of the ERK-1 and ERK-2 in Caco-2 (*dKO*) cells. **(E)** The ERK deficient *dKO* cells failed to reduce the paracellular flux of inulin upon starvation. **(F)** Co-Immunoprecipitation showed that in the absence of the ERK kinases, autophagy induction did not reduce the interaction of occludin with LAMP-2, beclin-1, and caveolin-1 in the *dKO* cells. The negative control includes immunoprecipitation with control IgG. **(G)** Quantification of beclin-1, caveolin-1, and LAMP-2 associated with occludin immunoprecipitates from panel F. Statistical analysis by one-way ANOVA with Tukey’s post-test. *, # and **, *p* < 0.05. **(H)** Starvation did not prevent the co-localization between occludin (green) and caveolin-1 (red) upon CSD treatment in the *dKO* cells. White bar = 5 μm. White arrows denote occludin-caveolin-1 co-localization. Statistical analysis by one-way ANOVA with Tukey’s post-test. *,*p* < 0.05; **, *p* < 0.01; ***, *p* < 0.001; ns – non-significant.

### Occludin protects against inflammation-induced barrier loss

We next assessed if the starvation-induced upregulation of occludin protects the epithelial TJ barrier against immune injury. Profuse production of IFN-γ, TNF-α and IL-17A by hyperactive gut resident lymphocytes, the consequent gut inflammation, and loss of the gut epithelial barrier are hallmarks of IBD.^32, 33^ We, thereby, assessed the effect of the three cytokines individually on the paracellular flux of inulin. Basolateral treatment of Caco-2 cell monolayers with either IFN-γ or TNF-α alone had minimal effect on the paracellular barrier while treatment with IL-17A produced a mild increase of paracellular inulin flux (**Fig. S5A, B, and C)**. In a potentiating effect to IL-17A induced barrier loss, a cytokine cocktail comprising all three cytokines had the maximum TJ barrier disruptive effect. The TER of Caco-2 monolayers incubated with the cytokine cocktail progressively decreased over time compared to control cells. However, when starvation-induced Caco-2 cells were incubated in the presence of the cytokines, the TER loss was prevented. Additionally, we observed that the flux of inulin was significantly increased upon cytokine treatment, which was prevented when the cytokine treatment was combined with starvation (**Fig. 5A and 5B**). We next assessed if exogenous occludin overexpression is protective against the cytokine-induced paracellular TJ barrier loss. For this, we performed a similar cytokine treatment on Caco-2 cells overexpressing occludin (**Fig 5C and D)** and observed no loss of the paracellular TJ barrier upon cytokine treatment (**Fig. 5E**) compared to empty vector control. Taken together these data highlight the physiologically relevant protective role of the increased levels of occludin protein during starvation-induced autophagy and the protective role of the occludin in maintaining the TJ barrier.

**Figure 5.**
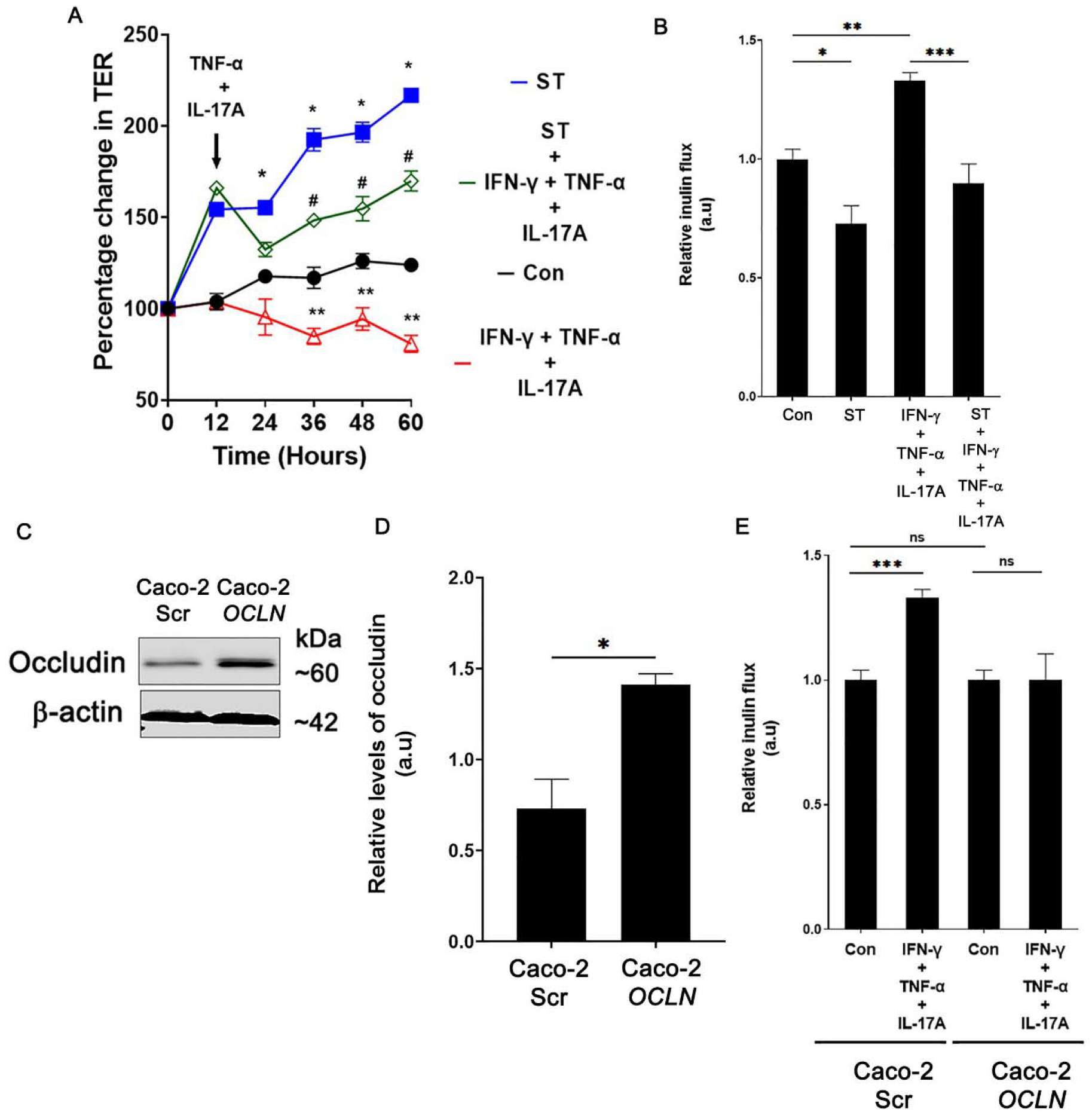
Autophagy-mediated and exogenous occludin overexpression protects against inflammation-induced barrier loss. **(A)** Starvation protected Caco-2 cell monolayers from IFN-γ + TNF-α + IL17A treatment-induced drop in TER. n = 4. Statistical analysis by paired Student’s *t-test.* *, *p* < 0.05 Con *vs.* ST; **, p < 0.05 Con *vs.* IFN-γ + TNF-α + IL-17A; #, p < 0.05 IFN-γ + TNF-α + IL-17A *vs.* ST + IFN-γ + TNF-α + IL-17A. Data represents of 4 independent experiments. **(B)** Starvation protected Caco-2 cell monolayers from IFN-γ + TNF-α + IL17A treatment-induced increase in paracellular inulin flux. n = 4. **(C)** Western blot showing baseline occludin levels in empty vector (GFP) and occludin overexpressing (GFP-*OCLN*) vector-transfected Caco-2 cells. **(D)** Quantification of occludin in panel C. n = 3 **(E)** Occludin overexpression protected against the IFN-γ + TNF-α + IL17A -induced disruption of the TJ barrier. Statistical analysis by Student’s *t-test* or one-way ANOVA with Tukey’s post-test. *, *p* < 0.05; **, *p* < 0.01; ***, *p* < 0.001; ns – non-significant.

### Autophagy induction protects against LPS and TNF-α induced intestinal injury in-vivo

Next, we examined the protective role of autophagy-induced increase in occludin levels *in vivo.* Lipopolysaccharides (LPS), major components of gram-negative bacterial cells walls are inflammatory antigens known to elicit intestinal injury and cause loss of occludin-associated TJ barrier.^34, 35^ Additionally, previous studies have also established the role of TNF-α as a pro-inflammatory cytokine that elicits intestinal injury and TJ barrier loss by promoting the endocytosis of occludin.^22, 36, 37^ Therefore, we used these antigens to study the role of autophagy-mediated TJ barrier enhancement. Intraperitoneal LPS injection in WT mice led to a significant loss in colonic occludin levels which was prevented upon autophagy-inducer rapamycin treatment (**Fig. 6A and B**). Rapamycin treatment successfully induced autophagy in mouse colonocytes as evidenced through degradation of the autophagy marker p62/SQSTM1. Loss of occludin levels was accompanied by increased intestinal permeability of inulin and rapamycin significantly protected against LPS-induced colonic (**Fig. 6C)** and small intestinal TJ barrier loss (**Fig. S6A)**. Rapamycin also prevented the LPS-induced increase in the association between occludin and caveolin-1 in mouse colonocytes (**Fig. 6D).** Similar to LPS, in the TNF-α injury model, intraperitoneal TNF-α administration increased the colonic (**Fig. 6E)** and small intestinal (**Fig. S6B)** permeability of inulin which was significantly attenuated by rapamycin. Rapamycin also prevented the TNFα induced increase in occludin caveolin-1 association in mouse colonocytes (**Fig. 6F).** To confirm the role of autophagy in the protective effect conferred by rapamycin seen in the LPS and TNF-α models described above, we performed the same experiments in autophagy-deficient a*tg*7 cKO mice. Acute deletion of *Atg7* in adult mice disrupted autophagy and reduced baseline occludin protein levels in the mice colon (**Fig. 6 G**). The a*tg*7 cKO mice also showed increased baseline colonic inulin flux compared to *Atg*7^fl/fl^ mice (**Fig. 6H)**. Moreover, unlike the *WT and Atg7*^fl/fl^ mice, rapamycin has no TJ barrier protective effect against LPS and TNF-α injury in these autophagy-deficient mice (**Fig. 6 I and J**). In a chemical colitis model, consistent with a previous report^29^, we have recently shown that a*tg*7 cKO mice have increased susceptibility to experimental dextran sodium sulfate (DSS) colitis which is reflected in higher disease activity index and severe colonic inflammation compared to control *Atg7*^fl/fl^ DSS mice.^11^ The a*tg*7 cKO DSS mice also showed remarkably increased colonic 4K dextran flux compared to *Atg*7^fl/fl^ DSS mice (**Fig. 6K**). Taken together, these studies provide evidence for the importance of the autophagy process and the TJ barrier enhancing role of occludin in-vivo and strongly underscore the role of autophagy-mediated increase in occludin levels against intestinal inflammation.

**Figure 6.**
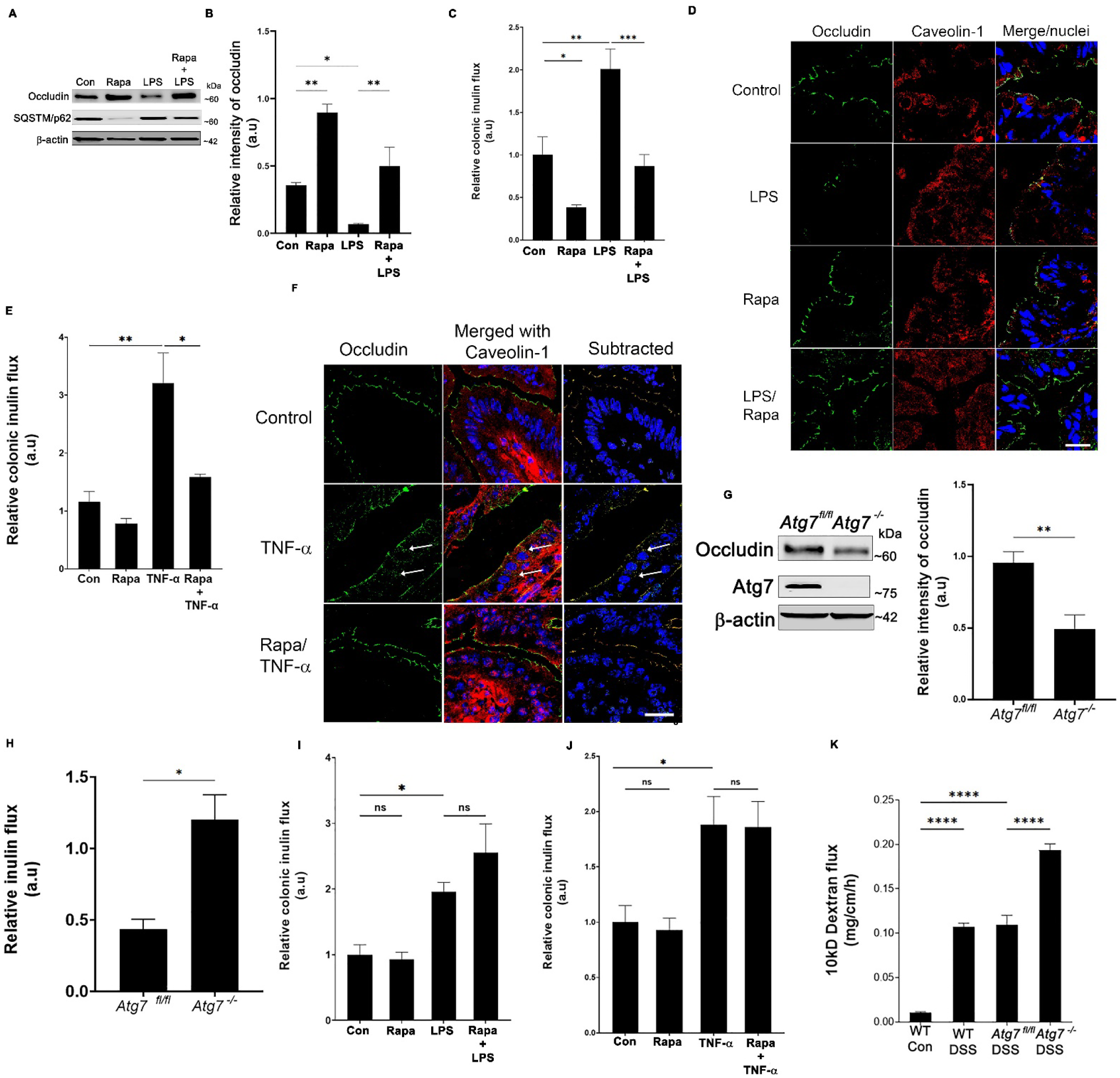
Autophagy protects against inflammation-induced TJ barrier loss *in vivo*. **(A)** Rapamycin treatment protected against LPS (0.2μg/kg body weight, 16 hours) induced reduction in occludin in mouse colonocytes. Degradation of SQSTM/p62 is shown as a marker of autophagy induction by rapamycin injection. **(B)** Quantification of occludin from panel A. n = 6/ group. **(C)** Rapamycin protected against LPS induced loss of colonic TJ barrier. n = 6/group. **(D)** LPS treatment increased the association of occludin (green) with caveolin-1 (red) which was prevented by rapamycin. White bar-10 μm. **(E)** Rapamycin protected against TNF-α (0.2μg/kg body weight, 4 hours)-induced loss of colonic TJ barrier. n = 6/group. **(F)** TNF-α also increased the association between occludin (green) and caveolin-1 (red) in WT mice colonic mucosa which was prevented upon rapamycin treatment. White bar-10 μm. White arrows show occludin – caveolin-1 co-localization (yellow). **(G)** Tamoxifen treatment of adult mice with Atg7 floxed and Ubc-CreERT2 alleles resulting in the loss of ATG7 protein in colonic mucosa (*Atg7*^-/-^ mice) compared to ATG7 floxed (*Atg*7^fl/fl^) control mice and caused a decrease in constitutive occludin levels in the colonic epithelial cells. Densitometry for occludin levels in *Atg*7^-/-^ mice. **(H)** The *Atg*7^-/-^ mice showed increased baseline colonic inulin flux compared to control *Atg*7^fl/fl^ mice. Rapamycin failed to prevent LPS (**I**) and TNF-α (**J**)-induced increase in colonic inulin flux in *Atg*7^-/-^ mice. **(K)** In the acute dextran sodium sulfate (DSS) colitis model, *Atg*7^-/-^ mice showed markedly increased colonic inulin flux compared to *Atg*7^fl/fl^ DSS and WT DSS mice. Statistical analysis by Student’s *t-test* or one-way ANOVA with Tukey’s post-test. *, *p* < 0.05; **, *p* < 0.01; ***, *p* < 0.001; ****, *p* < 0.0001; ns – non-significant.

### Autophagy upregulates occludin levels and enhances the TJ barrier in human colonic explants in an ERK1/2 dependent process

To assess the role of autophagy on occludin levels in the human colon, explant cultures from surgically resected diseased and adjoining normal colonic tissues from CD patients were treated with rapamycin. Both normal and diseased tissues incubated in rapamycin showed significantly increased levels of occludin compared to untreated tissues (**Fig. 7 A, B**). In addition to establishing the role of autophagy in occludin upregulation in the human colonic tissues, we further used this system to study the role of ERK1/2 in the autophagy induced upregulation of occludin and enhancement of the paracellular barrier. For this, we incubated surgically resected normal colonic tissues with rapamycin in the presence of the MAPK inhibitors U0126 or PD98059. Rapamycin treatment increased the levels of occludin in the tissues compared to the control, however, it was inhibited by U0126 (**Fig. 7 C, D**). Furthermore, rapamycin treatment also enhanced the TJ barrier of the human tissues as evidenced by the reduction of paracellular inulin flux which was abrogated by both U0126 and PD98059 treatment (**Fig. 7E).** Taken together these data show that autophagy induces upregulation of occludin in human colonic tissues and enhances the paracellular TJ barrier, in MAPK dependent manner.

**Figure 7.**
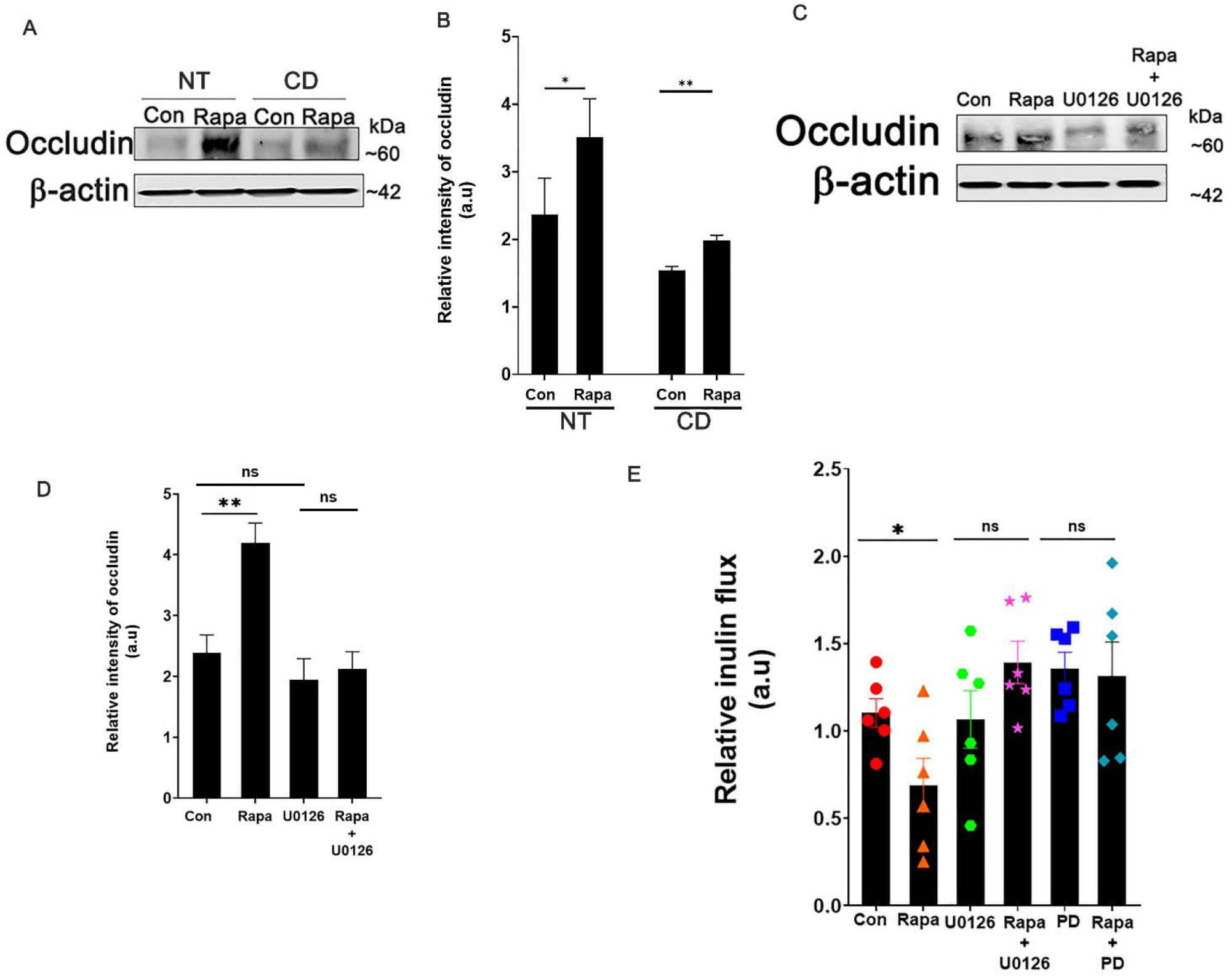
Autophagy enhances occludin levels in human colonic tissues in an ERK1/2 dependent mechanism. **(A)** Western blot for occludin from Rapamycin (300 nM) treated Normal (NT) and diseased (CD) mucosal tissue from patients diagnosed with Crohn’s disease. Rapamycin treatment significantly increased the levels of occludin in both NT and CD tissues. **(B)** Quantification of occludin levels in panel A. n = 5. **(C)** Western blot for occludin from normal human mucosa upon rapamycin treatment with or without ERK1/2 inhibitor U0126 (25 μM) Inhibition of ERK1/2 prevented the rapamycin-induced upregulation of occludin levels. **(D)** Quantification of occludin levels in panel C. N = 6/group. (**E)** Rapamycin treatment enhanced the TJ barrier and reduced the paracellular flux of inulin across human colonic tissues which were prevented upon inhibition of ERK1/2 with U0126 or PD98059 (20 μM). N = 6/ group. Statistical analysis by Student’s *t-test* or one-way ANOVA with Tukey’s post-test. *, *p* < 0.05; **, *p* < 0.01; ns – non-significant.

## Discussion

We have previously reported that nutrient starvation-induced autophagy reduces intestinal epithelial TJ permeability and enhances TJ barrier function via degradation of cation-selective, pore-forming TJ protein claudin-2 ^10, 11^. Here we show a novel role of the autophagy pathway, which is traditionally thought to be degradative, in selective preservation of the TJ protein occludin and reduction in the paracellular permeability of macromolecules. The starvation-mediated upregulation of the barrier-forming protein occludin contrasts its degradation promoting effect on the pore-forming protein claudin-2, and underscores two distinct mechanisms through which autophagy mediates the enhancement of the TJ barrier.

Our study shows that starvation induces occludin upregulation in different epithelial cell lines, mouse and human intestinal tissue, and tissue-derived organoids. We also showed that this upregulation of cellular occludin is not exclusive to starvation since this effect was also produced by several known pharmacological inducers of autophagy. Starvation increased Thr phosphorylation and localization of occludin to the TJs and resulted in the enhancement of the paracellular barrier against macromolecular flux. Pharmacological and genetic inhibition of the autophagy pathway prevented the starvation-induced TJ barrier enhancement, underscoring the role of autophagy in this process. We also highlight the importance of occludin in maintaining the paracellular barrier. Consistent with the previous reports ^38–41^, the baseline inulin flux in the cells lacking occludin (*OCLN*^-/-^) compared to control cells remained unaltered, however, autophagy-induced barrier enhancement was not observed in these cells. These observations agree with previous reports where mice lacking occludin show no morphological differences in the TJ architecture and intestinal TER compared to WT mice but show retarded post-natal development, age-dependent barrier dysfunction, and an increased propensity for gut injury indicating that occludin is crucial for TJ barrier maintenance. ^40,42–45^. Several studies, including our laboratory, have demonstrated that occludin is constitutively trafficked via caveolae.^11,20,46, 47,48^. Our present study demonstrates that autophagy enhances cellular occludin levels by reducing its endocytosis from the membrane and increasing its half-life. Furthermore, we show that autophagy shunts occludin off the constitutive degradation pathway by disrupting its association with caveolin-1 and Atg-6/beclin-1. We have previously reported that Atg-6/beclin-1 regulates constitutive degradation of occludin^23^, and our present findings suggest that engagement of beclin-1 in the autophagy pathway spares occludin from degradation.

The TJ barrier composition and integrity are regulated by different kinases and the protective role of the Ser/Thr kinases proteins ERK-1 and ERK-2 have been established previously.^49, 50^ Activation of ERK-1 and 2 by the AMP-activated protein kinase (AMPK) and subsequent activation of TSC2 or their AMPK independent activation by PKC^51^ is involved in promoting autophagy.^30, 52^ Our findings show increased phosphorylation and activation of the MAPK proteins ERK-1 and ERK-2 upon starvation-induced autophagy and pharmacological inactivation and genetic knockout of these kinases in Caco-2 cells nullified the starvation associated TJ barrier enhancement and occludin localization to the TJ barrier. Our data also underscores the importance of the ERK kinases in protecting occludin from degradation as, in their absence, autophagy induction failed to reduce the interaction of occludin with either caveolin-1, beclin-1, or LAMP2 or protect against CSD induced occludin endocytosis. Though we found that ERK1/2 does not have conventional caveolin binding motifs (CBMs), non-conventional CBMs in ERK1/2 are known to mediate interaction with the scaffold domain (SD) of caveolin-1. ^53^ This taken together with our observation that starvation enhances the association between the ERK kinases and caveolin-1 alongside the Caco-2 protein interactome upon starvation (data not shown), points to a possible mechanism of direct interaction between the ERK kinases and caveolin-1 which prevents the interaction between caveolin-1 and occludin.

To understand the relevance of autophagy-mediated occludin upregulation in inflammation, we assessed the role of autophagy and occludin in multiple *in-vitro* and *in-vivo* injury models of IFN-γ, TNF-α, IL-17A, LPS, and DSS colitis, which have previously been shown to increase the permeability of the gut epithelium.^16, 22, 34, 36, 37, 54^ We found that *in-vitro* autophagy induction by starvation successfully prevented the cytokine-induced increase in paracellular macromolecule permeability. Additionally, we show that occludin overexpression protects the Caco-2 monolayers from the cytokine-induced TJ barrier loss. This is unlike a previous report in MDCK II cells that shows depletion of the TJ barrier with overexpression of occludin during cytokine treatment.^48^ This difference is possibly due to the different cell line models or the ramification of autophagy induction in the regulation of intracellular pathways including caveolin-1. Furthermore, our *in-vivo* observations show that autophagy induction in the intestine of WT mice with rapamycin protects against the TJ barrier disruptive effect of LPS and TNF-α which is not recapitulated in the autophagy-deficient *Atg*7^-/-^ mouse intestine. Additionally, the overall severity of DSS colitis including the colonic paracellular macromolecule flux was markedly increased in *Atg*7^-/-^ mice. These observations along with the decreased baseline occludin expression and increased colonic macromolecule flux in *Atg*7^-/-^ mice further support our previous *in-vitro* findings^23^ that autophagy is essential for regulating constitutive occludin levels and the TJ barrier function.

In addition to the *in vitro* and *in vivo* approaches using epithelial cell lines and mouse models, we show that autophagy-induction increases occludin protein levels in healthy and IBD patient-derived colonic tissues. Our work on the human tissues also emphasizes the importance of ERK 1 and 2 proteins in the autophagy-induced phenotypes, since pharmacological inhibition of these kinases inhibited the TJ barrier protective role of autophagy in terms of both occludin levels and the paracellular inulin flux.

Presently, autophagy is known to degrade several membrane proteins including amyloid precursor protein (APP, a precursor for β-amyloid)^55^, Notch1^56^, focal adhesions^57^, and E-cadherin.^58^ Autophagy-mediated suppression of occludin degradation along with the fact that no other TJ barrier-forming proteins such as claudin-1, −3, −5 are altered during autophagy induction^10^, indicates the selective nature of autophagic occludin up-regulation. Occludin overexpression or promotion of occludin expression is generally known to restrict TJ barrier loss^22, 44, 59, 60^ but has also been shown to impact susceptibility for apoptosis under inflammatory conditions.^61^ We, however, did not observe increased apoptosis upon exogenous occludin overexpression in Caco-2 cells (data not shown). Several studies have shown that the inflamed intestinal mucosa in patients with active IBD has decreased occludin expression. ^62–64^ Thus, our findings of autophagy-induced upregulation of occludin are highly significant given the role of occludin in the TJ barrier along with the defects in the TJ barrier and autophagy reported in IBD. In conclusion, our present study demonstrates a unique function of autophagy in occludin up-regulation and enhancement of the TJ barrier. Autophagy may provide a novel tool to promote epithelial survival during intestinal inflammation.

## Abbreviations

ATG7: autophagy-related 7
Becn-1: Beclin-1
Cav-1: Caveolin-1
CBM: Caveolin Binding Motif
CSD: Caveolin-1 scaffolding domain peptide
Cas9: CRISPR associated protein 9
Con: control
Cyc: Cycloheximide
DSS: dextran sodium sulfate
ERK-1/2: Extracellular Signal Regulated Kinase 1 and 2
EBSS: Earle’s balanced salt solution
IBD: inflammatory bowel diseases
KD: knockdown
KO: knockout
LC3: Microtubule-associated proteins 1A/1B light chain 3B
mTOR: mechanistic target of rapamycin kinase
NT: Non-target
OCLN: occludin
phospho: Phosphorylated
Rapa: rapamycin
SQSTM1: sequestosome 1
ST: starvation
TER: Trans-epithelial resistance
Tyr: Tyrosine
Thr: Threonine
ULK1: unc-51 like autophagy activating kinase 1
WT: wild type.

## Funding

This research work was supported in part by the National Institute of Diabetes and Digestive and Kidney Diseases grant DK100562 (PN), DK114024 (PN), DK-106072 (TM), National Institute of General Medical Sciences grant P20-GM-121176, and Crohn’s & Colitis Foundation Award 694583 (MN). The content is solely the responsibility of the authors and does not necessarily represent the official views of the funding agencies. The authors also acknowledge support from the Peter and Marshia Carlino Fund for IBD Research.

## Acknowledgments

The authors thank the IBD and Colorectal Diseases Biobank, Transmission Electron Microscopy, Confocal Microscopy, and Animal Facility cores at the Penn State College of Medicine for their excellent technical assistance.

## Disclosures

No conflicts of interest, financial or otherwise, are declared by the author(s).

---

**Supplementary Figure 1:**

**A: Starvation increases the presence of occludin at the TJ.**

Caco-2 cells were starved for 18□h and subjected to immune-gold transmission electron microscopy using an anti-occludin antibody. The dense electron-dense (5 um) dark spots in the images represent occludin. The presence of occludin was found to increase in the highlighted TJ structure upon starvation. Scale bar = 400 nm.

**B:** Starvation enhances occludin levels in mouse colonids. Epithelial crypts containing stem cells were isolated from WT mice colons and grown into organoids *in-vitro.* Organoids were then differentiated into intestinal epithelial cells and subjected to starvation. Occludin levels were assessed by confocal-immunofluorescence microscopy. Starvation enhanced occludin levels on the apical membrane of the epithelial cells.

**C: Starvation alters the Threonine phosphorylation and reduces Tyrosine phosphorylation of Occludin.**

Caco-2 cells were starved for 24 hours followed by co-immunoprecipitation with an anti-occludin antibody. Occludin immunoprecipitates from control and starvation samples were then assessed for Tyrosine and Threonine phosphorylation by using anti-Tyrosine and anti-Threonine antibodies. Starvation increases the levels of phospho-Threonine occludin while reducing the levels of Tyrosine phosphorylation which enables membrane localization of the occludin protein.

**D: Claudin-2 knockout does not affect inulin flux.**

From our previous studies which assessed the effect of autophagy on claudin-2 degradation, we tested the effect of claudin-2 deletion on the baseline inulin flux and observed no significant difference between the baseline inulin flux of Scr and *CDN2*^-/-^ Caco-2 cells.

Statistical analysis by unpaired Student’s *t-test.* ns- non-significant.

**Supplementary Figure 2:**

**A: Starvation-induced upregulation of occludin is not caused by transcriptional changes.**

Caco-2 cells were incubated in EBSS for 24 hours and RNA was isolated from control and starved cells. We observed no significant change in the levels of occludin mRNA upon quantification by RT-qPCR between the control and starvation samples.

**B and C: Global caveolar endocytosis remains unaffected during starvation.**

Control and starvation-induced Caco-2 cell samples were used to assess levels of phospho-Y14-caveolin-1, a marker for caveolar endocytosis. Phospho-caveolin-1 levels increased upon starvation (B).

We assessed the effect of starvation on cholera toxin endocytosis (C). Control and pre-starved Caco-2 cells were incubated with cholera toxin for 30 minutes and 60 minutes at 37°C. We observed no significant difference in the ability of Caco-2 cells to endocytose cholera toxin upon starvation.

**Supplementary Figure 3: Effect of *ATG7* and *OCLN* knockout on Caco-2 cells.**

A. Subtracted confocal immunofluorescence image showing co-localization of occludin (green) with caveolin-1 (red) upon CSD treatment in *ATG7*^-/-^ Caco-2 cells. White bar – 5 μm, white arrows represent sites of occludin-caveolin-1 co-localization (yellow).

B. Fluorescence quantification of yellow color from Panel A.

C. CRISPR/Cas9 mediated deletion of occludin in Caco-2 cells did not affect the starvation-mediated enhancement of TER.

Statistical analysis by Student’s *t-test* or one-way ANOVA with Tukey’s post-test. *, *p* < 0.05; a, p < 0.05 Scr Con vs. ST; b, p < 0.05 *OCLN*^-/-^ Con vs ST.

**Supplementary Figure 4:**

**Inhibition of ERK1/2 prevents the starvation-induced increase of the membrane localization of occludin.**

**A.** Caco-2 cells were subjected to starvation and total protein was isolated. The protein samples were used to perform liquid chromatography followed by mass spectroscopy. The interactome of occludin protein was visualized using STRING.

**B.** Starvation increased the localization of occludin (green) to the membrane which is prevented by the ERK1/2 inhibitor U0126 (25 μM). Maximum intensity projection represented from 30 z-stacks of 0.30 μm. Representation of several fields from 3 separate Caco-2 monolayer samples. White bar = 10 μm.

**C.** Starvation significantly increases the TER of *dKO* cells. (* *p* < 0.01 *vs*. Scr con *vs*. ST; #*p* < 0.05 *dKO* con *vs*. ST).

**D.** Subtracted confocal immunofluorescence image showing co-localization of occludin with caveolin-1 upon CSD treatment in *dKO* Caco-2 cells. White bar – 5 μm, white arrows represent sites of occludin-caveolin-1 co-localization (yellow).

**E.** Fluorescence quantification of yellow color from Panel A.

Statistical analysis by paired or unpaired Student’s *t-test* test. *, *p* < 0.05 Scr Con *vs.* ST; #, *p* < 0.05 *dKO* Con *vs*. ST.

**Supplementary Figure 5: Effect of cytokines on the paracellular inulin flux.** Filter-grown Caco-2 cell monolayers were basolaterally treated with 20 ng/ml human IFN-γ **(A)** or 20 ng/ml human TNF-α **(B)** or 50 ng/ml human IL-17A **(C).** IL-17A produced a mild but significant increase in inulin flux. Statistical analysis by unpaired Student’s *t-test* test. *, *p* < 0.05; ns-non-significant.

S**upplementary Figure 6:**

Intraperitoneal rapamycin injection protected against LPS **(A)** and TNF-induced **(B)** barrier loss and prevented the increase of inulin flux in the small intestine of WT mice. Statistical analysis by one-way ANOVA with Tukey’s post-test. *, *p* < 0.05; **, p < 0.01; ns-non-significant.

## Notes

### Competing Interest Statement

The authors have declared no competing interest.

