## Supplementary figures and images for "Autophagy Increases Occludin Levels to Enhance Intestinal Paracellular Tight Junction Barrier"

### Supplemental Figure : 1

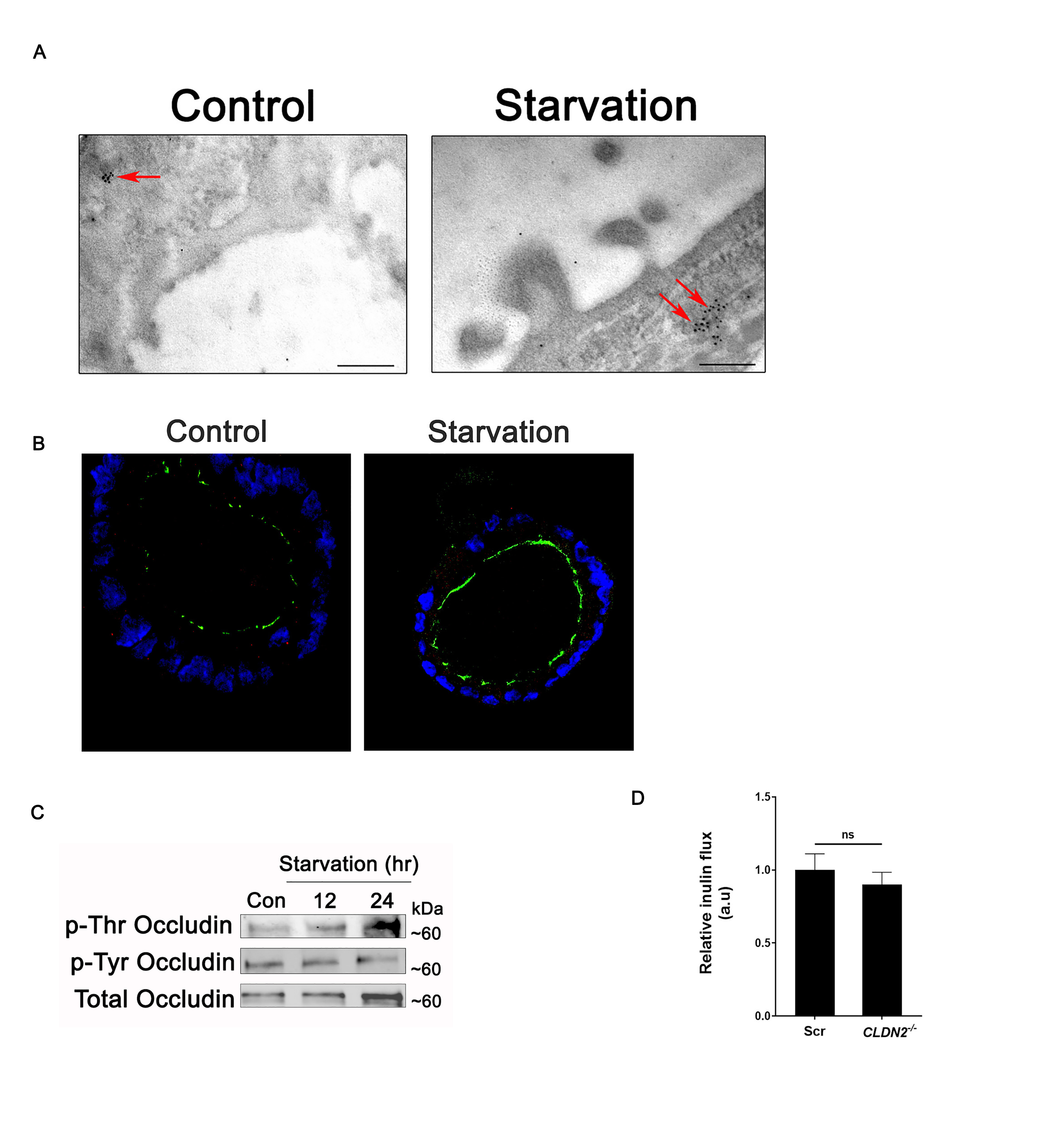

### Supplemental Figure: 2

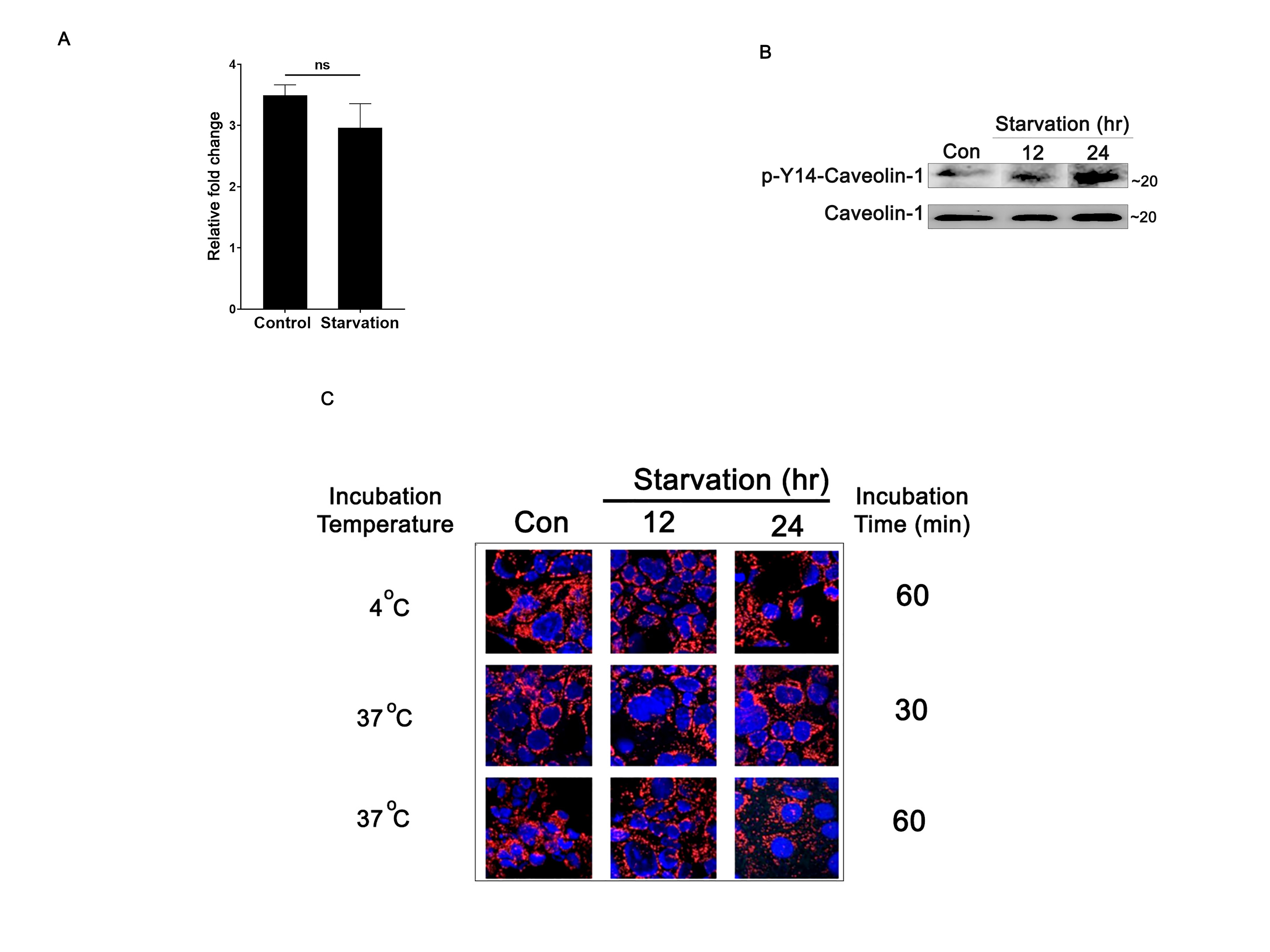

### Supplemental Figure: 3

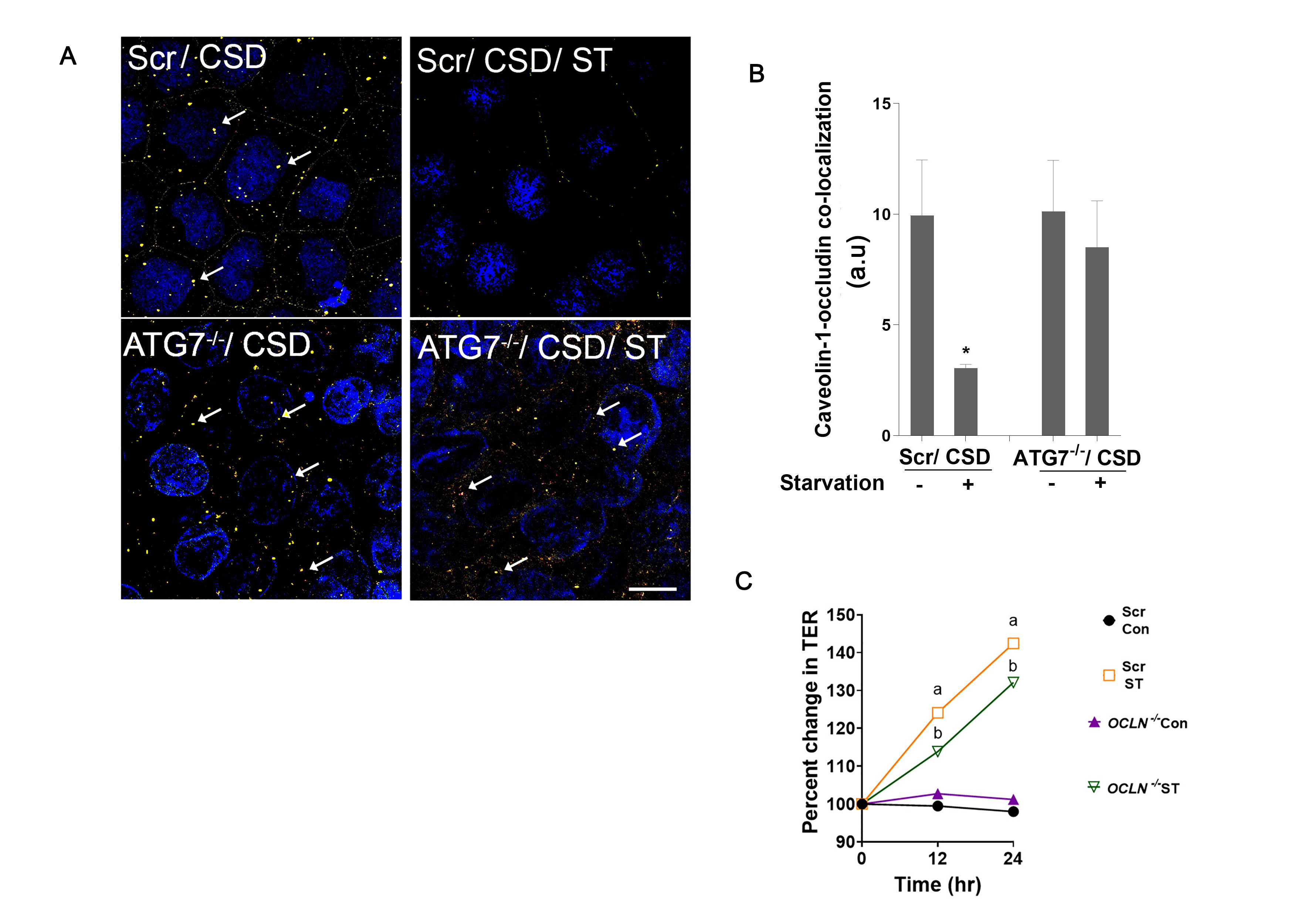

### Supplemental Figure: 4

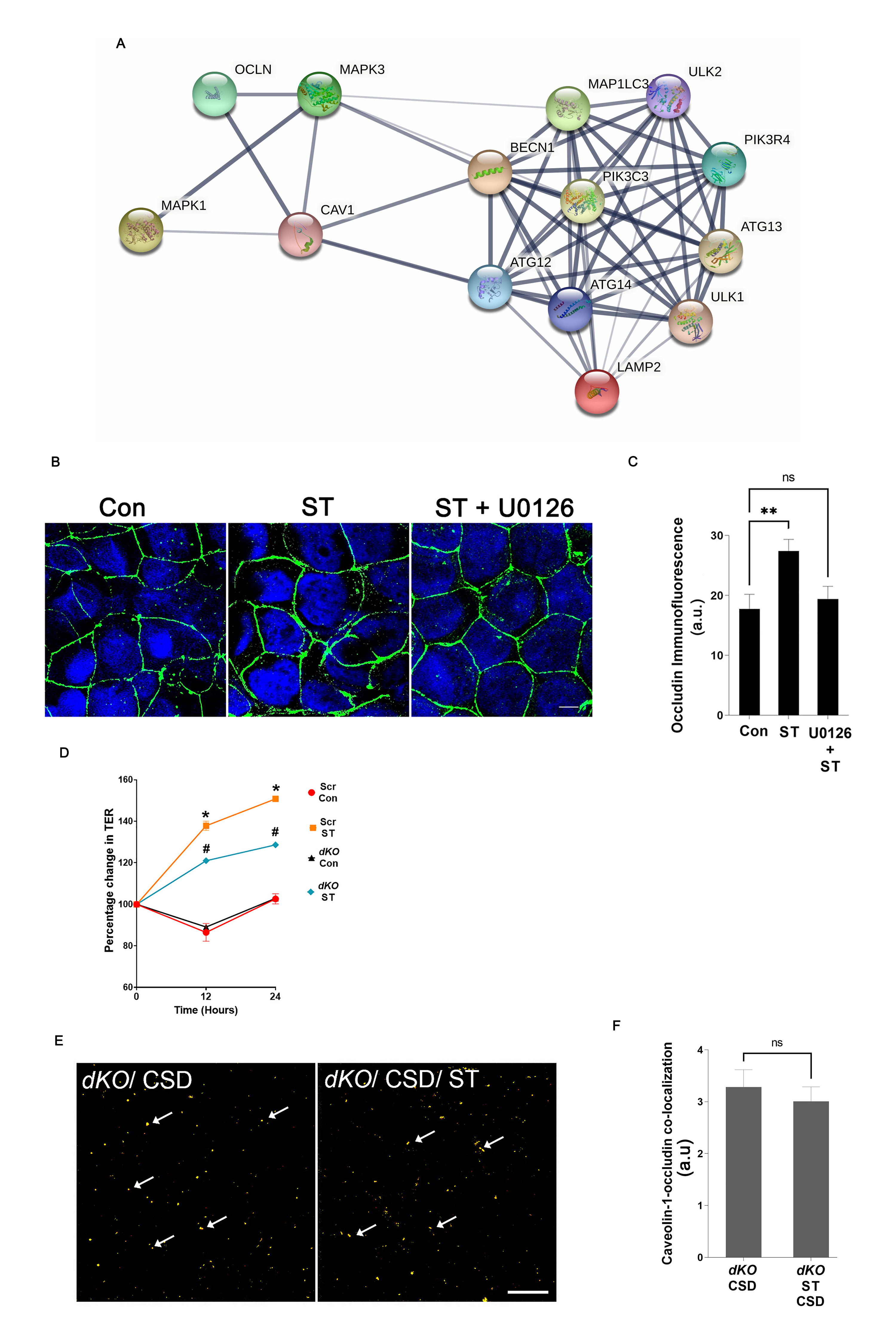

### Supplemental Figure: 5

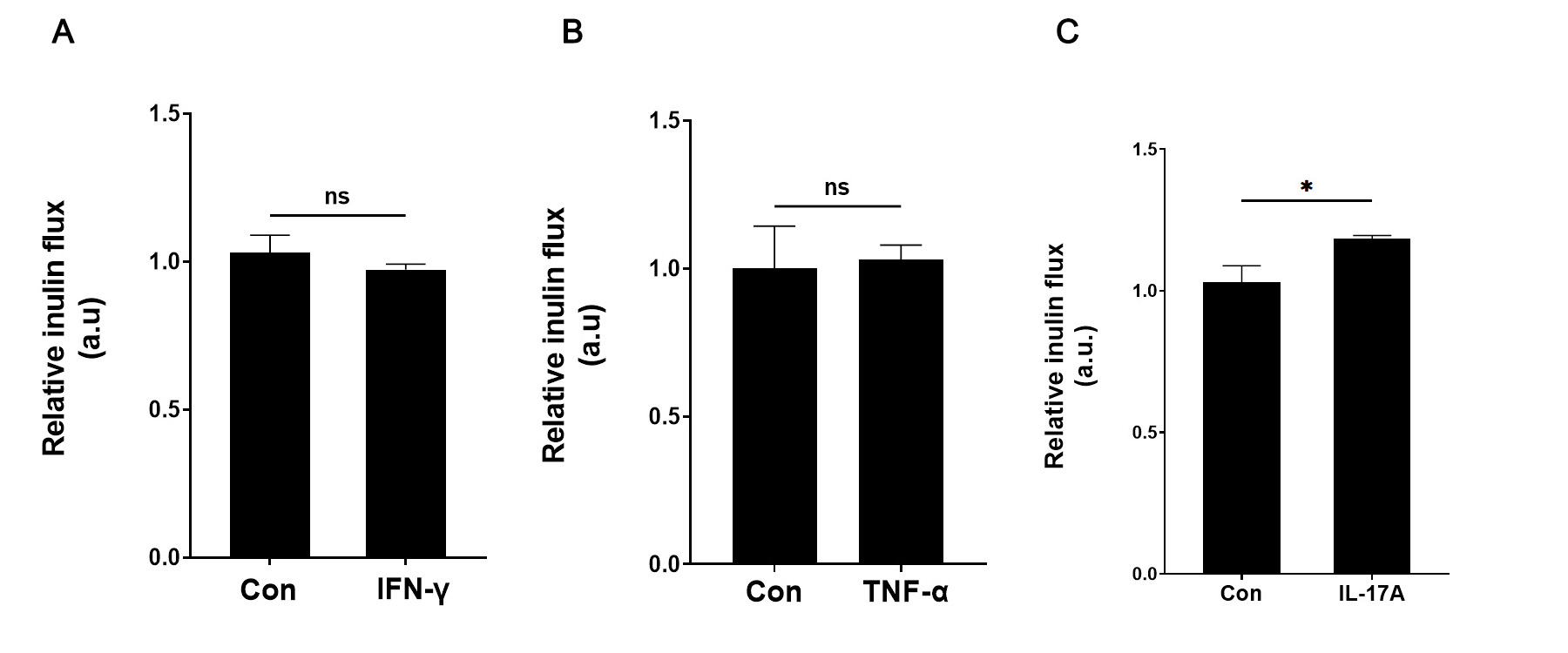

### Supplemental Figure: 6

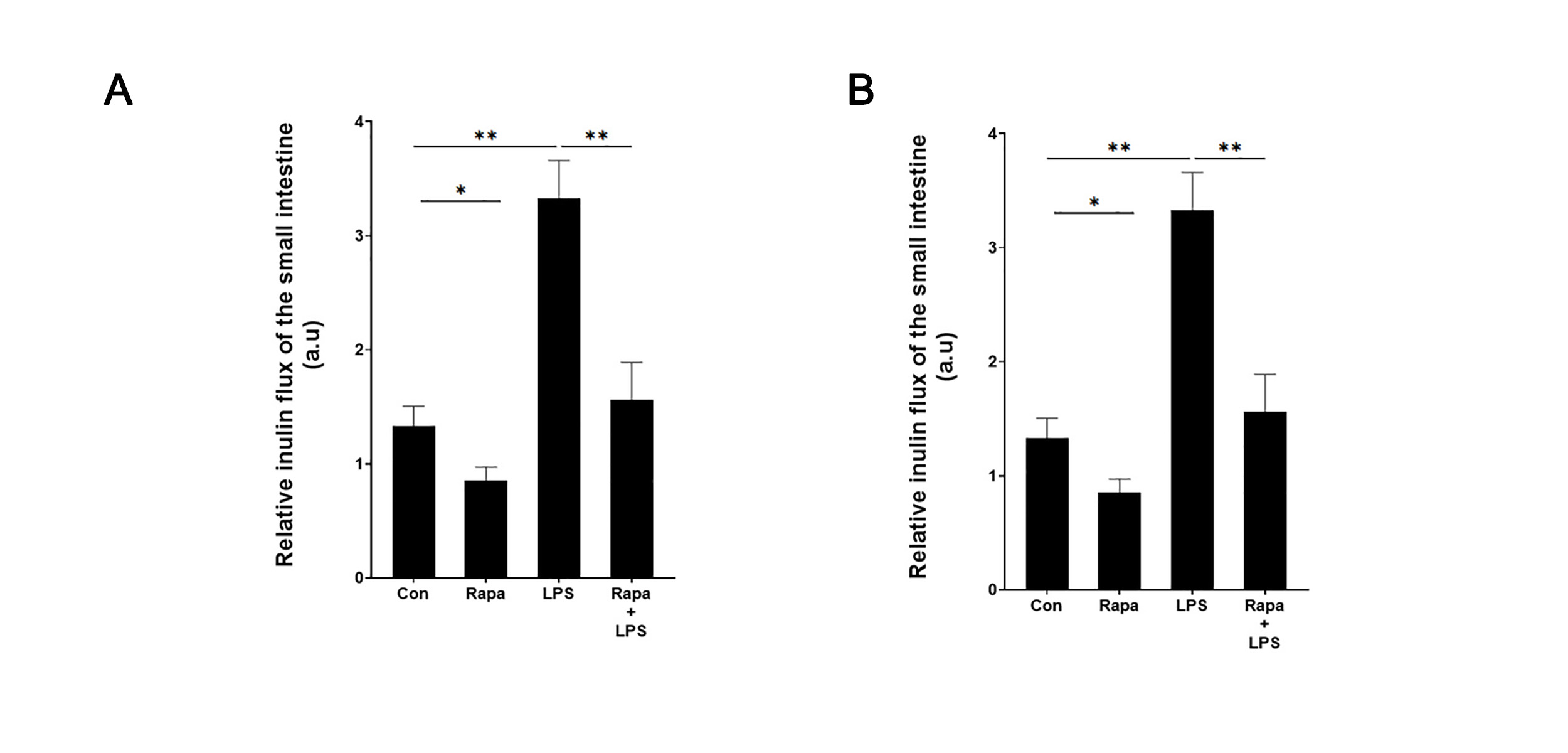
